# Vasculogenesis Potential of Mesenchymal and Endothelial Stem Cells Isolated from Various Human Tissues

**DOI:** 10.1101/049668

**Authors:** Rokhsareh Rohban, Nathalie Etchart, Thomas R. Pieber

## Abstract

Neo vessel formation can be initiated by co-transplantation of mesenchymal stem cells (MSC) with endothelial colony-forming cells (ECFC). The two adult stem cell types can be isolated and expanded from a variety of tissues to be used for regenerative applications pro-angiogenesis.

Here we performed a systematic study to evaluate the neo-vasculogenesis potential of MSC and ECFC isolated from various human tissues. MSC were isolated, purified and expanded *in vitro* from umbilical cord (UC) and umbilical cord blood (UCB), white adipose tissue (WAT), bone marrow (BM), and amniotic membrane of placenta (AMN).

ECFC were isolated from UC and UCB, WAT and peripheral blood (PB). ECFC and MSC and were co-transplanted admixed with extracellular matrix (Matrigel^®^) at a ratio of 5:1 to immune-deficient NSG mice, subcutaneously. The transplants were harvested after two weeks and the state of vessel formation and stability in the explants were investigated using immune-histochemical methods. The number of created micro-vessels was quantified using Hematoxylin & Eosin (H&E) staining followed by image J quantification.

Results showed that ECFC and MSC possess variable capacity in contributing to neo-vasculogenesis. WAT and UCB-derived ECFC and WAT, UCB and BM-derived MSC are most potent cells in terms of neo-vessel formation *in vivo*. UC-derived ECFC and AMN-derived MSC have been shown to be least potent in contributing to neo-vasculogenesis. This variability might be due to variable phenotypes, or different genetic profiles of MSC and ECFC isolated from different tissues and/or donors.

The findings might give an insight into better regenerative strategies for neo-vessel formation *in vivo*.

## 1. Introduction

Neo-vasculogenesis or development of novel micro-vessels by stem and progenitor cells is crucial in tissue engineering and regeneration as well as pathological aspects such as tumor growth as a result of pathological vessel development (Carmeliet and Jain, 2011). In case of neo-vessel formation, migration of transplanted and/or circulating endothelial progenitor cells results in forming outer layer of newly established micro-vessels. Mesenchymal stem cells or pericytes may also contribute in vessel formation by maintaining vasculature stability (Reinisch et al., 2007, Losordo and Dimmeler, 2004, Carmeliet and Jain, 2011).

In the process of embryonic generation, neo-vessel formation occurs according to a parallel process of formation of novel micro-vessels on one side and regression of other vessels on the other side (Risau and Flamme, 1995). The onset of neo-vessel formation is presumably through the cellular communication through secreted cellular factors that could serve as signaling mediators in various signaling cascades (Dasari et al., 2010, Dumont et al., 1992, Rohban et al., 2013). In this study, a well-established neo-vessel formation model (Au et al., 2008, Mead et al., 2008, Reinisch et al., 2009) has been used to test the hypothesis of whether the vasculogenesis potential of endothelial colony forming cells and mesenchymal stem cells is different depending on their tissue of origin. The findings can increase the knowledge on the vasculogenesis capacity of the progenitor cells that have been isolated from various human tissues.

This study shows that ECFC and MSC isolated from different human tissues possess seminal potential in terms of contribution to neo-vessel formation.The finding suggest that WAT-, UCB-and BM-derived ECFC and MSC are most potent in course of vessel formation *in vivo*, whereas UC-derived ECFC and AMN-derived MSC have been shown to be least potent in contributing to neo-vasculogenesis. This variability might be due to variable phenotypes leading to different genetic profiles of the cells isolated from various sites. The result of this study might be beneficial in establishing more effective regenerative strategies.

## 2. Materials and Methods

### 2.1. Ethics Statement

Human tissue samples were collected upon written informed consent from healthy individuals according to procedures approved by the Ethical Committee of the Medical University of Graz (Protocols 19-252 ex 07/08, 18-243 ex 06/07, 21.060 ex 09/10, 19-252 ex 07/08). Adult tissue samples were obtained after receiving written informed consent from healthy donors. Umbilical cord (UC) and umbilical cord blood (UCB) samples were collected following full-term deliveries with informed consent from the mothers according to the Declaration of Helsinki.

All animal experiments were carried out according to the 2010/63/EU guidance of the European Parliament on the welfare of laboratory animals. Protocols obtained the approval of the Animal Care and Use Committee of the Veterinary University of Vienna on behalf of the Austrian Ministry of Science and Research.

### 2.2. Preparation of ECFC and MSC Culture Medium

ECFC isolation and expansion was carried out using endothelial basal medium (EBM-2) supplemented with 100 μg/mL Streptomycin, 100 U/mL penicillin, 2 mM L-Glutamine (all Sigma) as well as manufacturer-provided aliquots containing vascular endothelial cell growth factor (VEGF), human epidermal growth factor (EGF), insulin-like growth factor-1 (IGF-1) and human basic fibroblast growth factor (bFGF). The medium was also supplemented with 10% pooled human platelet lysate (pHPL) replacing fetal bovine serum (FBS) as described previously (Reinisch et al., 2007, Schallmoser et al., 2007b, Reinisch et al., 2009, Hofmann et al., 2012b).

MSC isolation and expansion were performed using α-modified minimum essential medium (α-MEM, M4526; Sigma-Aldrich; St. Louis; MO, USA; www.sigmaaldrich.com) supplemented with 100 μg/mL Streptomycin and 100 U/mL Penicillin to minimize the bacterial contamination, 2 mM L-Glutamine (all Sigma;) and 10% pHPL (Schallmoser et al., 2007a, Bartmann et al., 2007, Schallmoser and Strunk, 2009). To avoid platelet lysate solidification in the medium, heparin (2 U/mL, Biochrom AG) was added to the α-MEM medium prior to pHPL supplementation.

### 2.3. ECFC and MSC Isolation and Expansion

ECFC were isolated and expanded from human umbilical cord, umbilical cord blood, adult peripheral blood and white adipose tissue (Prasain et al., 2012, Hofmann et al., 2009, Reinisch and Strunk, 2009, Mead et al., 2008, Szoke et al., 2012). For ECFC isolation from blood, cell culture was initiated no more than two hours after blood donation. To minimize cell loss, any unnecessary manipulation of freshly collected blood including red blood cell lysis or density gradient centrifugation was avoided. The heparin-containing blood was diluted in EGM-2 in a ratio of 1:4. The cultures were incubated at 37°C, 5% CO_2_ overnight and washed twice with pre-warmed PBS to remove non-adherent cells prior to addition of the fresh pre-warmed EGM-2 medium. The medium was replaced 2-3 times per week until the cobblestone-shaped ECFC colonies appeared (Reinisch et al., 2007, Schallmoser et al., 2007b, Reinisch et al., 2009, Hofmann et al., 2012b).

MSC were isolated and expanded from umbilical cord, umbilical cord blood, white adipose tissue, amnion and bone marrow (Shahdadfar et al., 2005, Szoke et al., 2012, Kita et al., 2010, Miao et al., 2006). Umbilical cord blood-derived mononuclear cells (UCB-MNC) were obtained from the interface using density gradient centrifugation (Ficoll-Paque^TM^-PLUS, StemCell Technologies Inc., Vancouver, Canada, www.stemcell.com). The mononuclear cells were washed with phosphate buffered saline (PBS) prior to be seeded in the culture flasks (Corning Inc., Acton, MA, USA) with a density of 1 x 10^6^ mononuclear cells /cm^2^ directly in supplemented α-MEM (Sigma). A complete medium replacement was carried out 2-3 days after seeding to promote the removal of non-adherent cells. The fresh medium was provided to the cultures twice per week. Mononuclear cell colonies consisting of approximately 50-80 cells in the primary cultures were subjected to trypsin (0.25% trypsin/1 mM ethylened iaminetetraacetic acid [EDTA], 1-5 min, 37°C; Sigma) after 2-3 weeks and were seeded into new flasks until reaching 80-90% confluence; thereafter, they were trypsinized and quantified prior to storage in liquid nitrogen. The cells were thawed for experimental assessment of expansion (Reinisch et al., 2007, Schallmoser et al., 2007b, Reinisch et al., 2009, Hofmann et al., 2012b).

Umbilical cord-derived MSC were isolated from Wharton’s jelly dissected from umbilical cord and cut into pieces (4-6 mm^2^). The tissue pieces were placed on the culture plate to allow for plastic attachment prior to addition of the culture medium (http://www.jove.com/author/AndreasReinisch). The cells were allowed to grow for 2-3 weeks.

In order to expand ECFC and MSC, the frozen cells of the first passage were thawed and seeded in 2,528 cm^2^ cell factories (CF-4, Thermo Fisher Scientific, Freemont, CA) using 500 mL EGM-2 and α-MEM medium, respectively. The cell-containing CF4s were maintained at 37°C, 5% CO_2_ and 95% humidity in an incubator. The cultures were subjected to medium replacement twice weekly to allow for the required nutrition accessibility and removal of the waste. Upon reaching 70-80% confluence, the cells were harvested using pre-warmed trypsin (50 mL per CF-4, 4-5 min., 37°C; Sigma Aldrich). After two washes with pre-warmed PBS and centrifugation (5 min, 300xg, 4°C), the harvested cells were quantified and checked for viability by means of Bürker Türk hemocytometer and trypan blue exclusion, respectively.

### 2.4. Flow Cytometric Characterization of ECFC and MSC

ECFC and MSC were stained with cell surface markers (Rohban et al., 2013, Reinisch et al., 2015) and the reactivity was evaluated using flow cytometry (Facs Calibur Flow cytometer, BD) according to the manufacturer’s instructions (Rohban et al., 2013, Reinisch et al., 2015). The ECFC and MSC cellular populations were checked for purity using marker profiling prior to use for *in vitro* and *in vivo* experiments (Rohban et al., 2013, Reinisch et al., 2015). Accordingly, trypsinized cell populations were considered to be pure when they were ≥95% positive for their positive markers and ≤2% positive for their negative markers (Dominici et al., 2006). ECFC and MSC populations were thereafter stored long term to be used for *in vitro* and *in vivo* assays.

### 2.5. Animal Experiments

Animal experiments in this project were performed following the 2010/63/EU guidance of the European Parliament on the welfare of laboratory animals. Protocols were approved by the Animal Care and Use Committee of the Veterinary University of Vienna on behalf of the Austrian Ministry of Science and Research. Immune-deficient NOD.Cg-Prkdc^scid^ Il2rg^tm1Wjl^/SzJ (NSG) mice were purchased from the Jackson laboratory (Bar Harbor, ME, USA), housed in specific pathogen-free (SPF) facility of the Medical University of Graz and were used for the *in vivo* experiments between 7 to 18 weeks of age. The immune-compromised NSG mice were used in order to avoid the immune response leading to the rejection of injected human cell transplants. ECFC/MSC were re-suspended with a ratio of 5:1 in ice-cold extracellular matrix (300 mL per plug, Cat. No. ECM 625, Millipore, Billerica, MA, USA) thereafter injected in form of subcutaneous plugs into the flank of the NSG mice (Melero-Martin et al., 2007, Au et al., 2008). Prior to injection, mice were anesthetized following the approval for animal handling (BMWF:-66.010/0082-II/10b/2009).

Mice were observed for the indicated time periods of two weeks after injection. At the defined time point after two weeks, the mice were sacrificed by cervical dislocation and the plugs were surgically dissected from the subcutaneous tissue, photographed using stereomicroscope (SZX12, Olympus) and fixed in neutral buffered 4% paraformaldehyde overnight to be used for histological assessments.

### 2.6. *In vitro* capillary-like structure formation

Umbilical cord blood, umbilical cord, white adipose tissue and peripheral blood-derived ECFC were seeded in 225 cm^2^ tissue culture flasks (BD Biosciences) and incubated in 37°C until confluence. Upon trypsinization followed by a washing step with pre-warmed PBS, 7.5 x 10^4^ ECFC were re-suspended in 2 mL EGM-2 supplemented with 10% pHPL and seeded on 9.2 cm^2^ polymerized Matrix (Angiogenesis assay kit; Millipore, Billerica, MA, USA) as described previously (Rohde et al., 2007). After 24 hours, endothelial networks were captured using a Color View III camera on an Olympus IX51 microscope with the analySIS B acquisition software (all Olympus, Hamburg, Germany). Numbers of branching points were evaluated using ImageJ software (http://imagej.nih.gov/ij/) as described (Hofmann et al., 2012a, Rohban et al., 2013).

### 2.7. Vessel Formation *In Vivo*

ECFC and MSC were subjected to an optimized isolation and purification procedure as previously indicated (Reinisch and Strunk, 2009) (see also Figure 1). ECFC were seeded in EGM-2 (Lonza) at a density of 1,000 cells/cm^2^ and MSC in α-MEM (Sigma-Aldrich, St. Louis, MO) at a density of 500 cells/cm^2^ in 2,528 cm^2^ cell factories (CF-4, Thermo Fisher Scientific, Freemont, CA).

**Fig.1.**
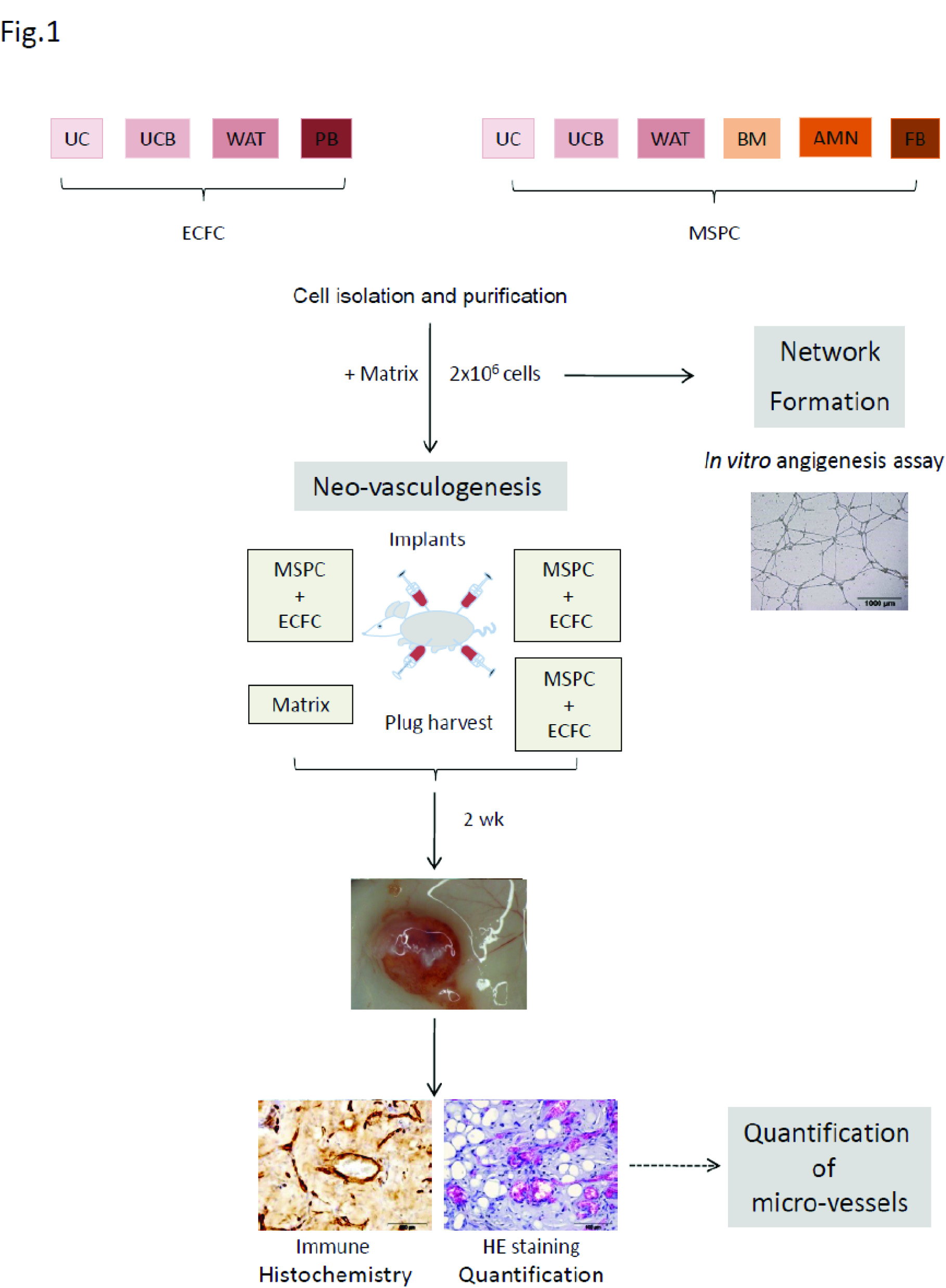

A total number of 2x10^6^ cells were used to investigate vessel formation. The combination of 6.4x10^6^ ECFC and 1.6 x 10^6^ MSC was re-suspended in 300 μL ice-cold extracellular matrix (*In vitro* angiogenesis assay kit, Millipore, Billerica, MA, USA) and injected subcutaneously into immune-deficient NSG mice (four plugs per mouse were injected, for every cell condition three mice were used). For every set of *in vivo* experiments, cell-free ‘Matrix only’ plugs (Millipore) were injected and used as controls (Figure 1).

The plugs were surgically removed after two weeks and were stored in Formaldehyde 3.7% to be used for histological experiments.

Mice were sacrificed at day 14 (two weeks) after implantation, and plugs were harvested from the subcutaneous sites (three mice and three plugs per cell combination per time point were used; Figure 2). Plugs explanted at two weeks (14 days) were used for histological experiments for confirmation of functional and stable vessel formation (Figure 2).

**Fig.2.**
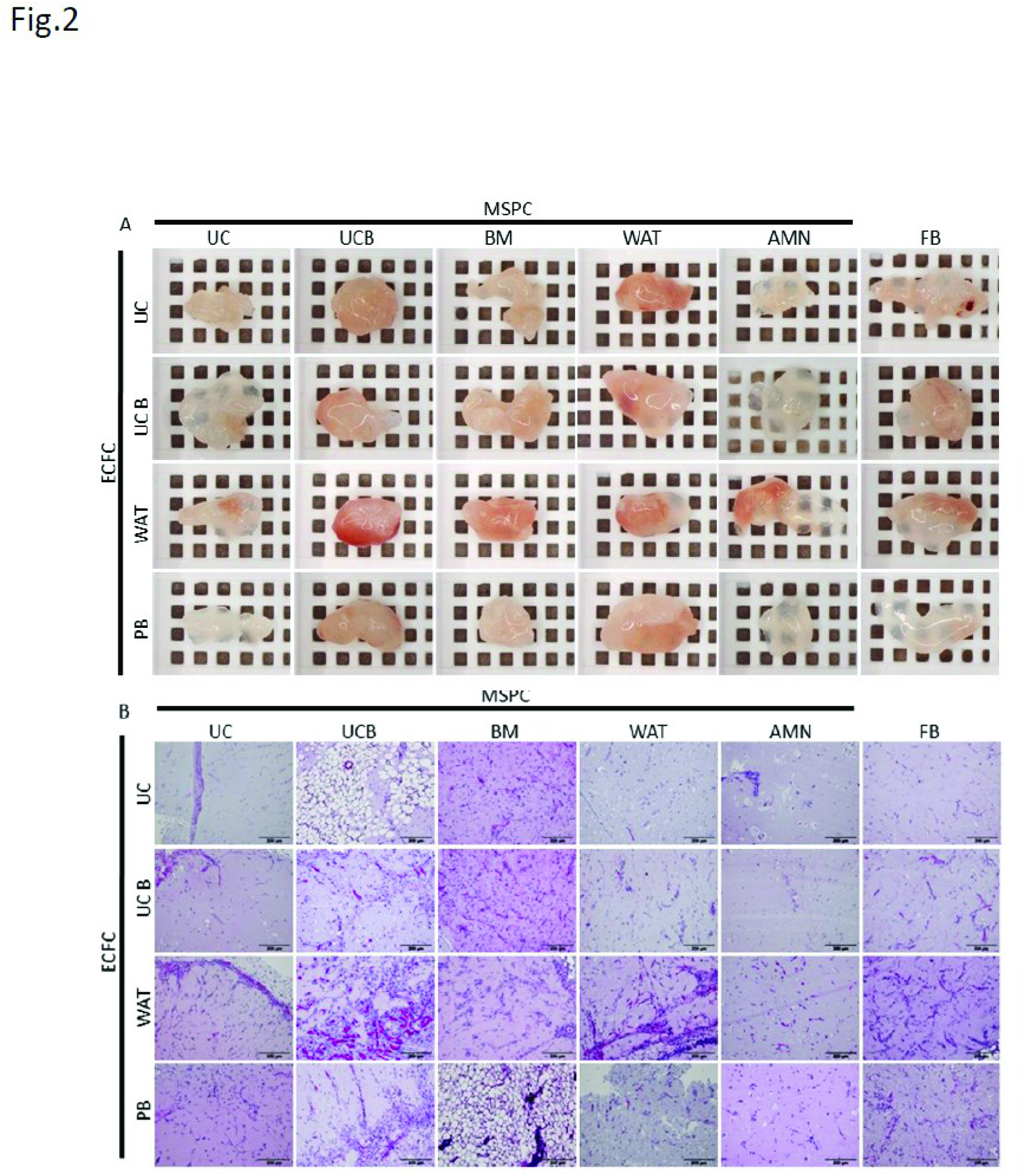

### 2.8. Histological Experiments and Analysis

Plugs harvested after two weeks were fixed using 3.7% neutral-buffered formalin in 4°C overnight thereafter dehydrated in a graded series of ethanol prior to being embedded in paraffin. Paraffin-embedded tissue sections (4μm) were de-paraffinized using xylene and descending alcohol series prior to Hematoxylin/Eosin (H&E), Movat’s pentachrome as well as immune histochemistry staining.

#### Hematoxylin-Eosin Staining (H&E)

Sections (4 μm) of formalin-fixed (3.7% neutral buffered, overnight) paraffin-embedded plugs were de-paraffinized using heat (68°C hot chamber) for an hour followed by treatment with xylene and descending alcohol series (5 min each) and stained by routine hematoxylin-eosin (H&E) protocol (Lillie, 1965). The sections were stained using Mayer’s hematoxylin (Sigma Aldrich) for 3 min, thereafter rinsed in running tap water and consequently stained with eosin phloxin (Sigma Aldrich) for 30 sec. Sections were then washed with running water, rehydrated in ascending alcohol series (2 min each) and mounted using TissueTek^®^ Glas ^TM^ mounting media (Sakura). The nuclei, cytoplasm and red blood cells were differentially stained blue, purple and red, respectively (Lillie, 1965). The protocol has been optimized based on the specimen properties.

To quantify created neo-vasculatures, luminal structures containing red blood cells were considered as perfused micro-vessels and counted by using ImageJ (http://rsbweb.nih.gov). For each condition, 5 high power fields (200x original magnification) of H&E-stained slides were examined by two independent observers using counting function in ImageJ software. For *in vivo* vasculogenesis experiments, micro-vessel density was quantified by analyzing the number of red blood cell-containing luminal structures.

#### Immune Histochemistry Staining

Sections (1.5 μm) of formalin-fixed (3.7% neutral buffered, overnight) paraffin-embedded plugs were de-waxed prior to antigen retrieval using heat (68°C/160 W, 40 min) at either pH 9 or pH 6, depending on the protein target and antibody properties followed by a descending alcohol series (2x xylol, 5 min each; 1x ethanol 100%, 5 min; 1x ethanol 90%, 5 min; 1x ethanol 70%, 5 min; 1x ethanol 50%, 5 min; 1x PBS, 5 min, distilled water, 5 min) as previously described (Hofmann et al., 2012a). Endogenous peroxidases were inhibited using H_2_O_2_ (15 min) and non-specific antibody binding was minimized using Ultra V Block (Thermo Scientific; 7-10 min), mouse-on-mouse blocking (MOM, Vector; 90 min) and serum-free protein inhibiting reagent (Dako; 30 min). Sections were incubated (45 min, RT) with unconjugated monoclonal mouse anti-human antibodies against CD31 (clone: JC70A, 5.15 μg/mL, Dako), CD90 (clone: EPR3132, Abcam, Cambridge, MA, USA) Vimentin (clone: V9, 0.78 μg/mL, Dako), von Willbrand factor (vWF, clone F8/86, Dako) or appropriate amount of IgG1 (BD) as the control. The signal was developed using Ultravision LP large volume detection system horseradish peroxidase (HRP) polymer (Thermo Scientific) followed by diaminobenzidine (DAB) or alkaline phosphatase detection system using either fast blue or fast red (Vector) according to the manufacturer’s protocols. Avidin-biotin inhibiting (Vector) was used before staining with biotinylated polyclonal rabbit anti-human CD90 (EPR3132, Abcam) and biotinylated monoclonal goat anti-rabbit IgG1 (BD). Streptavidin-horse radish peroxidase conjugate (Dako) and diaminobenzidine (DAB) were used to detect and visualize the positive signal. Sections were counter-stained using hematoxylin for 1-2 min. The protocol has been optimized based on the specimen and antibodies properties.

In order to quantify vasculogenesis in ECFC+MSC explants or plugs, luminal structures filled with RBC representing hu-CD31 and vimentin positive signal were considered as perfused functional vessels and quantified by two independent observers using five high power microscopic fields (200x original magnification) of H&E-stained sections from the related plugs using ImageJ software. The number of perfused neo-vessels was quantified using ImageJ software (http://rsbweb.nih.gov).

#### Movat’s Penta-chrome Staining

Paraffinized sections (4 μm) were hydrated using distilled water and stained in alcian blue (stain for ground substance and mucin, Dako, 20 min). The slides were washed under running water for 10 min followed by alkaline alcohol treatment (to convert the alcian blue into insoluble monastral fast blue) for 1-2h and washing steps (10 min) to remove alkaline alcohol. The sections were incubated with Verheoff’s hematoxylin solution (15 min, Sigma Aldrich) to promote nuclei and elastic fiber staining. Upon rinsing with dH_2_O (4 times, each time for 2 min), the slides were exposed to 2% aqueous ferric chloride several times to promote differential staining of the different tissues. The differentiation process was checked microscopically for black elastic fiber staining versus gray background. Upon washing steps using sodium thiosulfate (1 min) and distilled water (5 min), the sections were stained with scarlet-acid fuchsine (8:2, Dako) for 90 sec (Stain for fibrinoid, fibrin-intense red, muscle-red). After rinsing steps in distilled water and 0.5% acetic acid water, the slides were incubated with 5% aqueous phosphotungstic acid (5-10 min) and checked microscopically for further differentiation to collagen indicator (pale pink) versus bluish ground substance. After differentiation was completed, the slides were rinsed in 0.5% acetic acid water (5 min, 2X) followed by 100% alcohol (5 min, 3X).

### 2.9. Statistics

All data are shown as mean ± standard deviation (SD). Statistical differences were determined using unpaired student’s t-test or ANOVA using post-hoc Bonferroni test for multiple comparisons. Differences were considered as significant (*) when the P-value was less than 0.05, as very significant (**) when P-value was less than 0.001 and as extremely significant (***) when P-value was less than 0.0001.

## 3. Results

### 3.1. ECFC and MSC possess variable capacity in contributing to neo-vasculogenesis.

Subcutaneous transplantation of ECFC+MSC isolated from different human tissues in an well-established ratio of 5:1 (Reinisch et al., 2009) was carried out to perform a comparative study on neo-vasculogenesis potential of these mesodermal progenitor cell types *in vivo* (Figure 1).

Plugs containing ECFC which were isolated from UC, UCB, WAT and peripheral blood (PB) and MSC which were isolated and propagated from umbilical cord (UC), umbilical cord blood (UCB), white adipose tissue (WAT), bone marrow (BM), amniotic membrane of placenta (AMN), and mesenchymal-like cells (fiobroblast (FB)) were transplanted with the ratio of 5:1 as pre-vasculogenesis transplantation model as previously described (Hofmann et al., 2012a, Rohban et al., 2013). The use of different sources for isolation of ECFC and MSC was to compare the impact of the cellular contribution to neo-vessel formation with regards to the tissue of origin.

Vessel formation was evaluated macroscopically and histologically in the ECFC+MSC explants containing 2x10^6^ cells (three mice and three implants per condition and time point were used; Figure 2). Macroscopic features of the explanted plugs indicated the development of perfused red blood cell-containing vessels in the plugs *in vivo* that was further confirmed by immune histochemistry analysis. The red color of the plugs explanted after two weeks, macroscopically indicated patent red blood cell-containing micro-vessel formation after ECFC+MSC (Figure 2).

The result confirmed the ability of ECFC/MSC to develop micro-vessels regardless of the source of isolation. However, a significant difference in the number of created micro-vessels is detectable (Figure 3), which emphasizes different vasculogenesis potential of ECFC and MSC and with regards to the tissue of isolation/origin.

**Fig.3.**
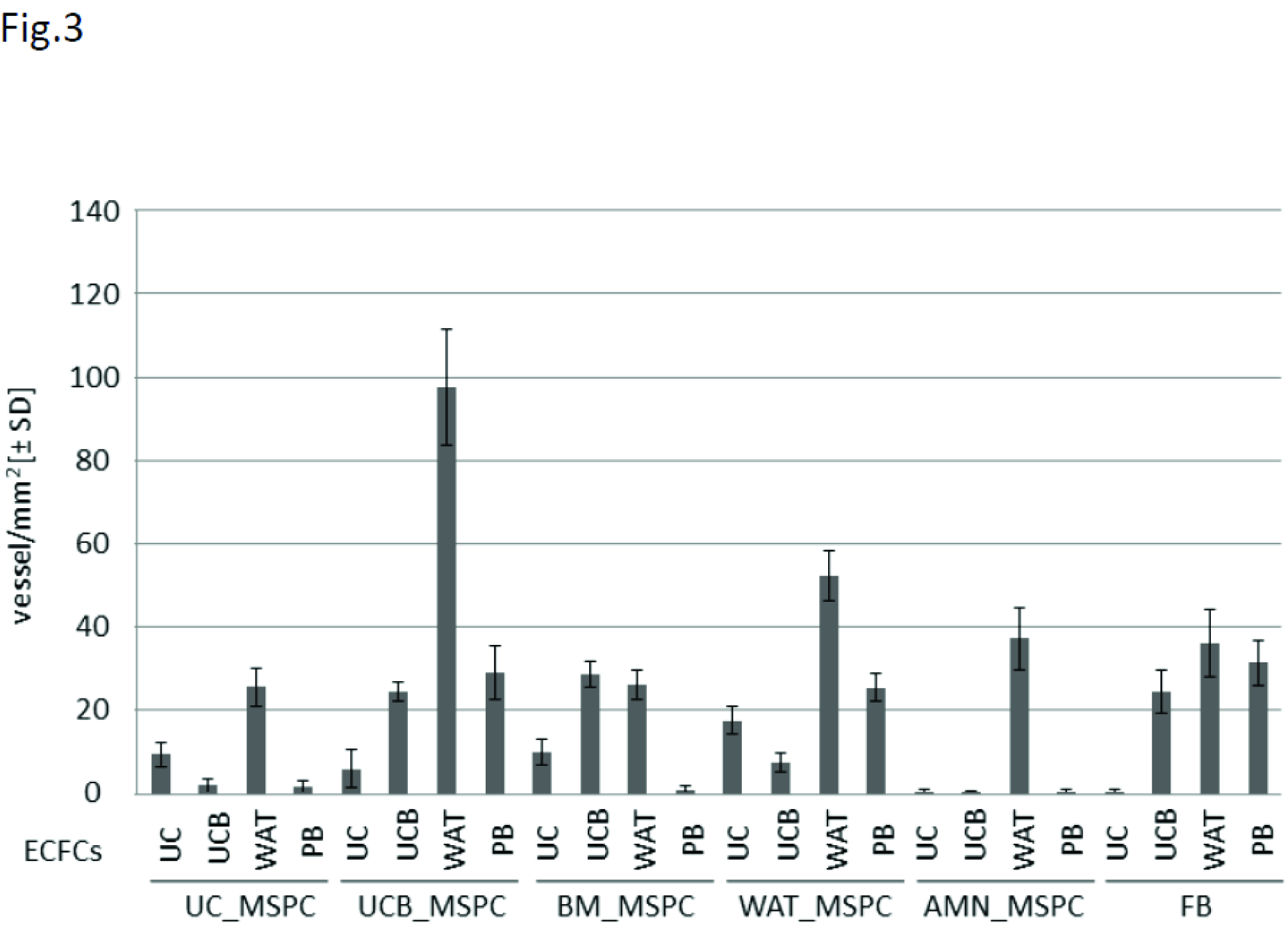

Human vimentin immunehistochemistry staining indicates that the established micro-vessels are originated from human mesodermal derived stem and progenitor cells (Rohban et. al, 2013).

### 3.2. ECFC isolated from various tissues possess different number of branching points in terms of capillary-like network formation *in vitro*.

ECFC isolated from UC and UCB, WAT and PB were used to perform in vitro angiogenesis assay. The results showed various numbers of branching points for ECFC isolated from different tissues. WAT-ECFC and UCB-ECFC showed a higher number of branching points, whereas UC-ECFC established a lower number of branching points *in vitro* (Figure 4 under development…).

### 3.3. WAT and UCB-derived ECFC and WAT, UCB and BM-derived MSC are most potent in course of vessel formation *in vivo*.

The number of created neo-vessels has been quantified by using H&E staining and ImageJ quantification method according to established criteria (guideline): 1) a perfused vessel is a vessel that contains ≥ 2 red blood cells, and 2) has a distinct outer layer which is CD31 and/or vimentin positive.

The results showed that the number of developed neo-vessels is variable according to the injected cell combination. The most potent cell combinations with regards to the number of created micro-vessels *in vivo* are UCB-MSC/WAT-ECFC and WAT-ECFC/WAT-MSC (Figure 3).

### 3.4. UC-derived ECFC and AMN-derived MSC have been shown to be least potent in contributing to vessel formation.

The result of micro-vessel quantification data showed that the number of established micro-vessels in transplants containing UC-derived ECFC and AMN-derived MSC is of a lower quantity regardless of the other cell type admixed (Figure 3,5).

**Fig.5.**
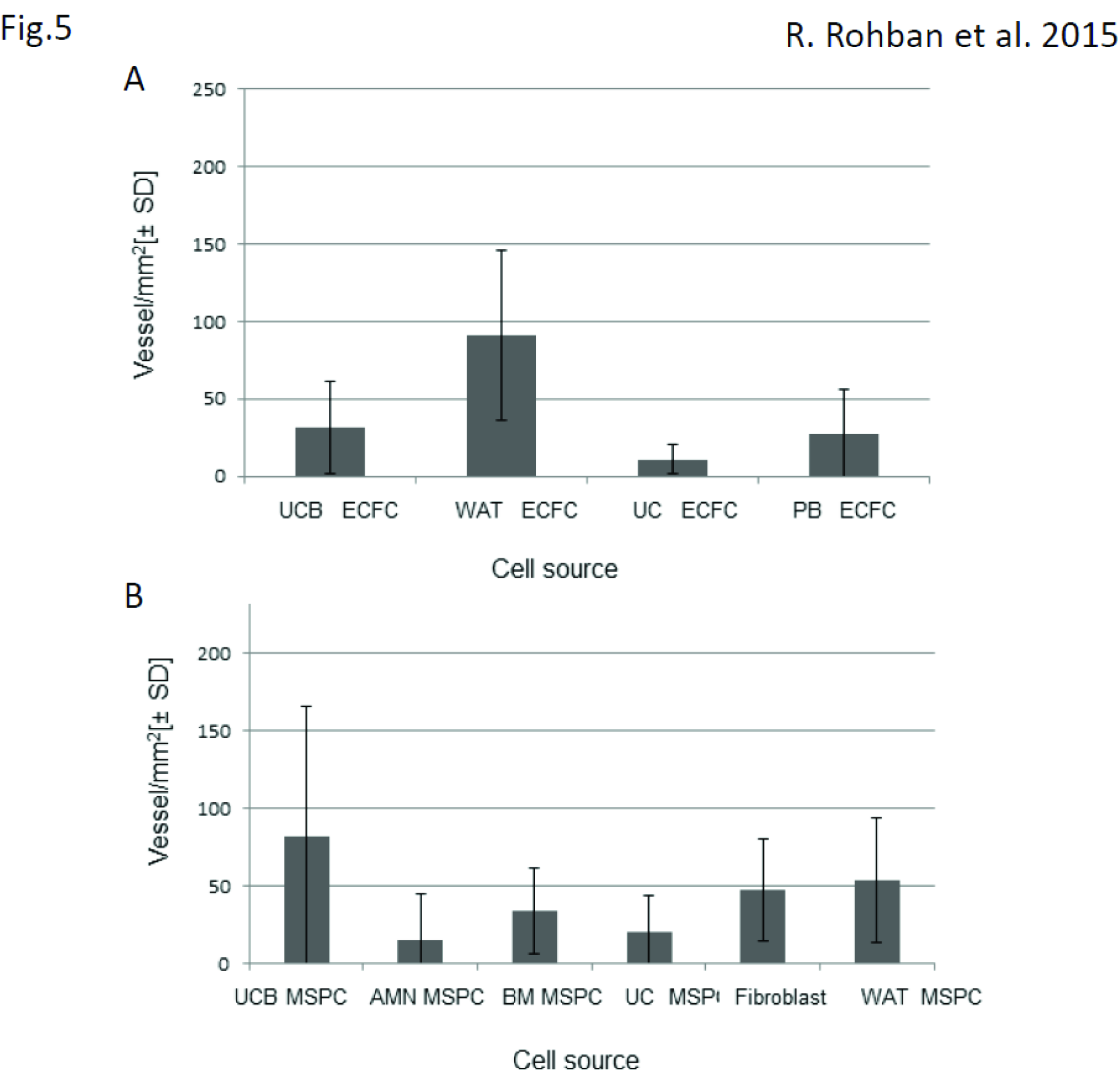

The neo-vasculogenesis capacity of the ECFC and MSC were also evaluated regardless of the other cell type admixed. Results showed that WAT-ECFC and UCB-MSC were most potent in contributing to neo-vessel establishment, whereas UC-ECFC and AMN-MSC were shown to be least potent in contributing to neo-vasculogenesis *in vivo* (Figure 5 A, B).

## Discussion

De novo vessel formation (neo-vasculogenesis) is an essential step in organ regeneration, wound healing, inflammation as well as tumor growth (Segura et al., 2002, Elmore, 2007, Krysko and Vandenabeele, 2008). Vascular structure formation occurs upon migration and replication of endothelial colony forming cells (ECFC) as the backbone of newly formed vessel (Segura et al., 2002, Elmore, 2007, Krysko and Vandenabeele, 2008). Mesenchymal stem cells (MSC) as pericytes also serve as vessel supporters and maintain micro-vessel stability (Reinisch et al., 2007, Schallmoser et al., 2007b, Reinisch et al., 2009, Hofmann et al., 2012b).

The aim of this study was to investigate the potentially existing variability in neo-vasculogenesis capacity amongst ECFC and MSC, isolated from different human tissues, and to identify the most or least vasculogenesis-competent cells with regards to their tissues of origin. The capacity of these cells in contributing to capillary like network formation *in vitro* was also investigated as a pre-indicator for the vasculogenesis potential of cells *in vivo*.

The aim was accomplished by 1) isolation, purification and expansion of ECFC and MSC from 6 different human tissues including UC, UCB, BM, AMN, PB, WAT, and by 2) investigation of the vasculogenesis capacity of isolated MSC and ECFC in contributing to vessel formation and 3) determination of the most competent source from which the high potential, vasculogenesis-competent cells can be isolated.

The number of created micro-vessels was quantified based on established criteria for identification of perfused neo-vessels, and Image J was used to quantify the perfused micro-vasculature to allow for comparison of the number of created micro-vessels in every transplant with different cell combination. This strategy allowed us to identify the most potent tissue from which the ECFC and MSC can be isolated and can be used for neo-vasculogenesis application in regenerative medicine.

In a previous study, we showed that caspase and kinase signaling pathways play a crucial role in neo-vasculogenesis *in vivo* in the transplants containing UC-ECFC/UC-MSC (Rohban et al., 2013). Whether the same signaling molecules are involved in neo-vasculogenesis, when cell types from other sources are included, remains unclear and will be subjected to further studies.

In the previous study, the vessel formation potential of three different cell combinations within transplants including UC-ECFC/UC-MSC, WAT-ECFC/UCB-MSC and UCB-ECFC/BM-MSC has been investigated in order to assess the impact of caspase blockers in neo-vessel formation *in vivo* (Rohban et al., 2013). The results showed that the number of neo-vessels in non-treated and also caspase inhibitor-treated WAT-ECFC/UCB-MSC transplants was significantly higher compared to the other two cell combinations (Rohban et al., 2013). It can therefore be concluded that WAT-ECFC enhance the vasculogenesis potential even when combined with the least potent MSC source e.g. AMN; Figure 3).

The results showed that the vessel formation potential of Fibroblast (FB) was not significantly different with UCB-MSC in contributing to neo-vasculogenesis when admixed with PB-ECFC and/or UCB-ECFC, whereas WAT-ECFC enhanced the vasculogenesis potential of Fibroblast as well as other tested MSC sources.

The variability of the vasculogenesis potential amongst the stem and progenitor cells isolated from various tissues might be due to variable phenotypes, or different genetic profiles of the differentially located cells. There could be, however, similarity in genetic profiles of the cells isolated from close and related origins e.g. UC, UCB, AMN. The findings might give an insight into better regenerative strategies for vessel regeneration *in vivo*.

